# Antibiotic pressures in livestock and meat systems: evidence and interventions for a One Health future

**DOI:** 10.1101/2025.09.03.673933

**Authors:** Yi Lu, Hongyue Ma

## Abstract

Antibiotic use (ABU) in livestock is a major driver of antimicrobial resistance, yet global levels, determinants, and long-term trajectories remain poorly resolved. We compiled a harmonized dataset across countries for 2020 and a longitudinal series of meat production from 1961 to 2023 to quantify patterns, evaluate socioeconomic and regulatory correlates, and forecast to 2050 using a production linked proxy model. Total ABU correlated strongly with national meat output (Spearman r = 0.906, *p* < 0.001). ABU intensity in milligrams per kilogram meat was highly heterogeneous and showed no global association with production. Gross domestic product per capita was unrelated to total use or intensity, learning adjusted years of schooling related only weakly to intensity, and legal controls on use or sales alone did not improve efficiency. Projections indicate a near linear rise of about 1,663 tonnes per year, reaching 131,828 tonnes in 2030 and 165,094 tonnes in 2050, with cumulative use in 2020 to 2050 surpassing 1950 to 2020. Reducing intensity by ten to fifty percent by 2050 would avert 16,509 to 82,548 tonnes. Findings motivate an integrated roadmap of source control, farm and market stewardship, and scalable surveillance using production as a proxy within a One Health framework.

**Highlights:**

1. Modern livestock production remains heavily dependent on antibiotics.
2. Higher antibiotic use in livestock correlates with lower usage efficiency at the global scale.
3. Current legislative controls on antibiotic use have failed to enhance usage efficiency.
4. Since the 1950s, global antibiotic use in livestock has followed a steady linear rise, projected to continue through 2050.

## 1. Introduction

Global meat consumption has significantly increased in recent decades, driven primarily by rapid population growth, urbanization, and rising incomes^1^. From 1961 to 2023, global per capita meat consumption nearly doubled, surging from 22.95 kg/year to 41.96 kg/year, while total meat production expanded approximately fivefold, from 71.36 million tonnes to 340.29 million tonnes^2^. To meet this escalating demand, livestock production systems have undergone substantial intensification and industrialization, characterized by high-density animal farming practices^3,4^. While these intensified practices have facilitated higher productivity, they have concurrently increased the risk of infectious disease outbreaks, necessitating routine use of antibiotics for disease prevention, growth promotion, and treatment^5,6^.

The widespread and intensive use of antibiotics in animal agriculture is increasingly recognized as a critical public health and environmental concern. Agricultural antibiotic residues can disperse through manure and wastewater, contaminating soil, groundwater, and surface waters, creating reservoirs for antimicrobial resistance (AMR)^7–9^. Additionally, residues in animal-derived food products directly expose human populations, potentially facilitating the emergence and dissemination of resistant pathogens, which compromise therapeutic efficacy in human medicine^10^. Consequently, livestock operations themselves become significant evolutionary incubators for antibiotic-resistant bacteria, perpetuating a cycle of escalating antibiotic usage and resistance proliferation^11^.

Compared to human medical applications, antibiotic use (ABU) in agriculture is often subject to less stringent regulatory oversight, with lower thresholds for prescription, easier access, and less transparent monitoring and reporting systems^12^. These regulatory shortcomings are particularly pronounced in low- and middle-income countries (LMICs), where livestock production is often economically critical, and awareness and education regarding antibiotic stewardship remain insufficient^13,14^. For instance, antibiotic usage rates vary dramatically worldwide in 2020: France reported an agricultural ABU of 22 mg per population-corrected unit (PCU), whereas this figure was 208 mg/PCU in China, 253 mg/PCU in Mongolia, and as high as 338 mg/PCU in Thailand^15^. Such disparities highlight both the substantial potential and urgent need for improving ABU efficiency globally.

Despite heightened global attention and regulatory efforts to curb agricultural antibiotic overuse, substantial variations persist in the effectiveness of these interventions^16^. In many regions, existing regulatory frameworks fail to achieve meaningful reductions in antibiotic consumption or improvements in usage efficiency^17,18^. Furthermore, policies and market-based interventions often inadequately address fundamental drivers of antibiotic misuse, including inadequate surveillance systems, economic incentives, and limited awareness among stakeholders^16^.

To date, comprehensive analyses of global ABU patterns in livestock, and their relationships with meat production, regulatory effectiveness, and socio-economic factors remain scarce. To address this critical gap, our study systematically investigates global patterns of agricultural ABU, identifies key determinants influencing antibiotic consumption and efficiency, evaluates the effectiveness of current regulatory measures, and projects future trends from 1950 to 2050. Based on these insights, we propose a novel policy framework focusing on precision antibiotic stewardship and stringent source-level regulation, aiming to balance the global demand for animal protein production with the imperative of mitigating AMR risks.

## 2. Methods

### 2.1 Data sources and selection

Data used in this study were derived from publicly available datasets provided by Our World in Data (OWID), specifically selected for their robustness, standardization, and global representativeness. The following key variables for the year 2020 were included: ABU in livestock (measured in tonnes/year) by countries or regions for the year 2020 (https://ourworldindata.org/grapher/antibiotic-use-livestock-tonnes); ABU per kilogram of meat in (ABU/kg meat) livestock (mg/kg meat, standardized by Population-Corrected Units, PCU) for 2020 (https://ourworldindata.org/grapher/antibiotic-usage-in-livestock); total meat production (tonnes) data for 2020, encompassing cattle, poultry, sheep/mutton, goats, pig meat, and wild game (https://ourworldindata.org/grapher/meat-production-tonnes); specific livestock meat production (tonnes) data (pig meat, beef and buffalo meat, sheep and goat meat, poultry meat) for 2020 (https://ourworldindata.org/grapher/global-meat-production-by-livestock-type); meat supply per capita (kg/person/year) for 2020 (https://ourworldindata.org/grapher/meat-supply-per-person); per capita consumption (kg/person/year) by specific meat types (pig meat, beef and buffalo meat, sheep and goat meat, poultry meat) for 2020 (https://ourworldindata.org/grapher/per-capita-meat-consumption-by-type-kilograms-per-year); regulatory status indicating whether countries or regions had enacted laws prohibiting antibiotic growth promotion without prior risk analysis for 2024 (https://ourworldindata.org/grapher/regulations-antibiotics-growth-promotion); countries or regions’ legislative status on antimicrobial sales for terrestrial livestock for 2024 (https://ourworldindata.org/grapher/laws-antimicrobials-livestock); Gross Domestic Product per capita (GDP per capita, PPP-adjusted international dollars, constant 2017 values) for 2020 (https://ourworldindata.org/grapher/gdp-per-capita-maddison-project-database); and average learning-adjusted years of schooling (LAYs), integrating both education quantity and quality for 2020 (https://ourworldindata.org/grapher/learning-adjusted-years-of-school-lays). Additional data on ABU/kg meat in livestock (mg/kg meat, standardized by Population-Corrected Units, PCU) from 31 European countries or regions spanning 2010-2022 (https://ourworldindata.org/grapher/antibiotic-use-in-livestock-in-europe), along with contemporaneous meat production, consumption, GDP per capita, and average LAYs were also retrieved from OWID.

Data screening was performed to ensure reliability and completeness. For analyses involving total ABU and ABU/kg meat in livestock, 181 countries or regions with complete datasets were retained. Similarly, 181 countries or regions were selected for analyses involving meat production and consumption due to consistent data availability. For analyses of GDP per capita and average LAYs, 148 countries or regions were selected due to varying data completeness. For regulatory analyses, 171 countries or regions were included.

### 2.2 Data analysis

#### 2.2.1 Global antibiotic usage and efficiency

Given the substantial variation and skewness in total ABU and ABU/kg meat in livestock, log_10_^(n+1)^ transformations were applied to normalize the distribution for statistical robustness. Spearman correlation analyses were conducted using SPSS software (IBM SPSS Statistics 28.0) to assess relationships between total ABU (log-transformed tonnes) and ABU/kg meat in livestock (log-transformed mg/PCU) across 181 countries or regions for 2020. Scatterplots, heatmaps, and regression analyses were produced using GraphPad Prism 10 (GraphPad Software, San Diego, CA, USA).

Spearman correlation analyses were also performed for 181 countries or regions to examine relationships between ABU in livestock and total meat production, meat consumption, and specific meat-type production and consumption, employing log_10_^(n+1)^ transformed data. Similarly, correlations between GDP per capita, average LAYs, and ABU in livestock and meat production-consumption metrics were assessed using log-transformed data for 148 countries or regions.

#### 2.2.2 Global regulatory analysis of antibiotic usage

Spatial distribution maps illustrating the regulatory status regarding ABU in livestock for growth promotion and antimicrobial sales regulations were constructed for 171 countries or regions. Comparative analyses of total ABU in livestock, ABU/kg meat in livestock, total meat production, meat consumption per capita, GDP per capita, and average LAYs were performed between countries or regions with and without regulatory measures.

#### 2.2.3 Continental and European-specific analyses

Global data were further segmented into continental groupings (Africa, Asia, Europe, North America, South America, Oceania) to conduct continent-specific Spearman correlation analyses on the same variables as described above. All analyses and graphical representations were generated using SPSS and GraphPad Prism 10.

Line charts depicting the ABU/kg meat changes in livestock from 2010 to 2022 for 31 European countries or regions were generated using GraphPad Prism 10. Corresponding annual data for total meat production, total meat consumption, and GDP per capita were also visualized using GraphPad Prism 10. Total ABU in livestock from 2010 to 2022 was estimated based on ABU/kg meat in livestock and total meat production. The trends in ABU/kg meat in livestock, total ABU in livestock, total meat production, meat consumption per capita, and GDP per capita across the 31 countries or regions were categorized into three types: “Increase”, “Decrease”, and “Fluctuate”. The relative proportions of countries or regions within each trend category were visualized using pie charts.

#### 2.2.4 Predictive modelling of global antibiotic usage (1950-2050)

To estimate historical and future trends in antibiotic consumption in the livestock sector, we developed a predictive modelling framework integrating longitudinal meat production data with empirically derived antibiotic usage coefficients. Annual total meat production data from 1961 to 2023 were retrieved from the OWID database, covering all major livestock species and global regions. Linear regression models were fitted using GraphPad Prism 10 to capture temporal trends in global meat production and extrapolate projections from 1950 through 2050.

To estimate global antibiotic usage in livestock, we applied a production-linked coefficient derived from 2020 global data, quantifying the relationship between total ABU and total meat output. This coefficient was used to construct a secondary model predicting total antibiotic consumption based on projected meat production trajectories.

Country-specific models were also constructed for major livestock producers including China, the United States, Brazil, Canada, and Australia, using national meat production data combined with a globally derived antibiotic usage coefficient. Furthermore, intervention scenarios simulating a 10%, 30%, or 50% reduction in the annual growth rate of total ABU were implemented to assess their projected impact on future global antibiotic consumption. All model outputs were visualized using area plots to illustrate observed historical data, business-as-usual projections, and potential reductions under enhanced stewardship scenarios.

## 3. Results

### 3.1 Global landscape of ABU in livestock

Substantial disparities exist in the total ABU in livestock across countries and regions. In 2020, national-level ABU ranged from as low as 1.00 tonne to as high as 32,776.30 tonnes per year. The largest contributors were China (32,776.30 tonnes), Brazil (10,007.1 tonnes), the United States (5,747.7 tonnes), India (5,234.2 tonnes), and Australia (3,609.5 tonnes). Collectively, these five countries accounted for over half (57.84%) of the total ABU across the 181 countries or regions analyzed. Notably, China alone represented 33.04% of global reported use, followed by Brazil (10.09%), the United States (5.79%), India (5.28%), and Australia (3.64%) (Figure 1a). Consistent with these estimates, the global distribution map of total ABU highlights concentrated usage in major livestock-producing countries, particularly in East Asia, South Asia, and the Americas, whereas total usage levels are markedly lower in most African countries or regions and parts of Europe (Figure 1b). ABU intensity, ABU/kg meat in livestock, also exhibited pronounced global variation, ranging from 4 mg to 26,759 mg per PCU in 2020. High-use countries included Mongolia (253 mg/PCU), China (208 mg/PCU), Australia (165 mg/PCU), India (114 mg/PCU), and Canada (60 mg/PCU). In contrast, markedly lower values were observed in many European and African countries, including Sweden (5 mg/PCU), the United Kingdom (16 mg/PCU), France (22 mg/PCU), Kenya (25 mg/PCU), Nigeria (25 mg/PCU), and Algeria (33 mg/PCU). These low intensities may reflect either efficient and regulated use, particularly in high-income settings such as Europe, or limited access to antibiotics and low-intensity livestock production systems, as may be the case in parts of sub-Saharan Africa (Figure 1c).

**Figure 1.**
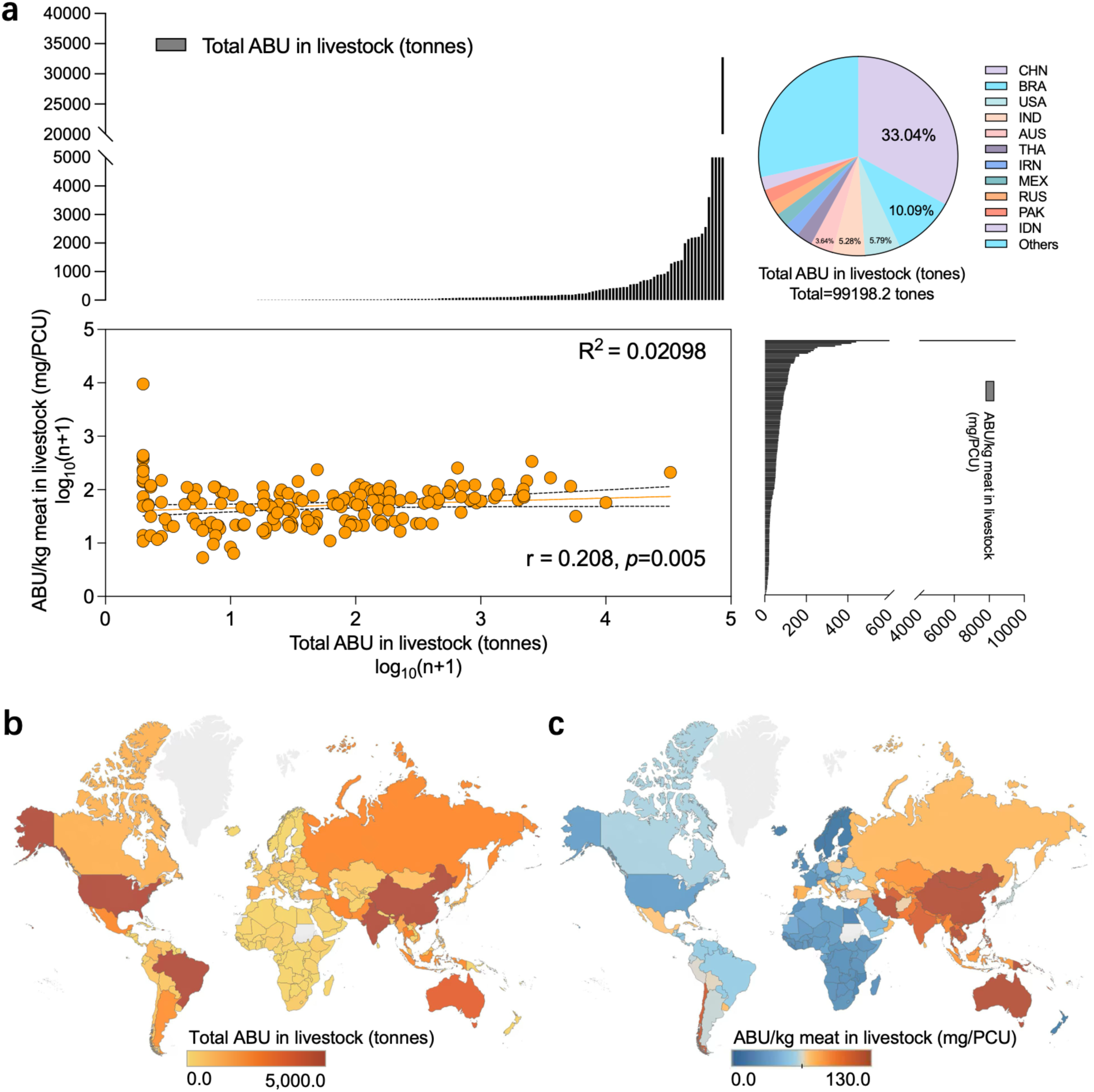
Global patterns and disparities in ABU in livestock. (a) Distribution and relationship between total ABU (tonnes, log_10_^(n+1)^-transformed) and ABU/kg meat (mg/PCU, log_10_^(n+1)^-transformed) in livestock across 181 countries or regions in 2020. Pie chart depicts national contributions to global total ABU. Bar plots show rank-ordered national totals for ABU and ABU/kg meat. (b) Geographic distribution of total ABU (tonnes) in livestock across 181 countries or regions in 2020. (c) Geographic distribution of ABU/kg meat (mg/PCU) in livestock across 181 countries or regions in 2020.

Importantly, a weak but statistically significant positive correlation was observed between total ABU in livestock (tonnes) and ABU/kg meat (mg/PCU) across countries or regions (r = 0.208, *p* = 0.005). This indicates that settings with higher absolute consumption tend to use more antibiotics per unit of production, reflecting lower use efficiency rather than gains from scale (Figure 1a). Taken together, these results indicate a decoupling between national consumption volume and ABU efficiency: higher volumes do not purchase efficiency gains and may coincide with more intensive unit use.

### 3.2 Global livestock ABU in relation to meat production and consumption

Total ABU in livestock was strongly correlated with total meat production across countries (r = 0.906, *p* < 0.001). This positive relationship remained consistent when disaggregated by meat type, including pig meat (r = 0.493, *p* < 0.001), beef and buffalo meat (r = 0.849, *p* < 0.001), sheep and goat meat (r = 0.651, *p* < 0.001), and poultry meat (r = 0.742, *p* < 0.001) (Figure 2a; Table 1). These findings emphasize the extensive reliance of meat production systems on antibiotic inputs.

**Figure 2.**
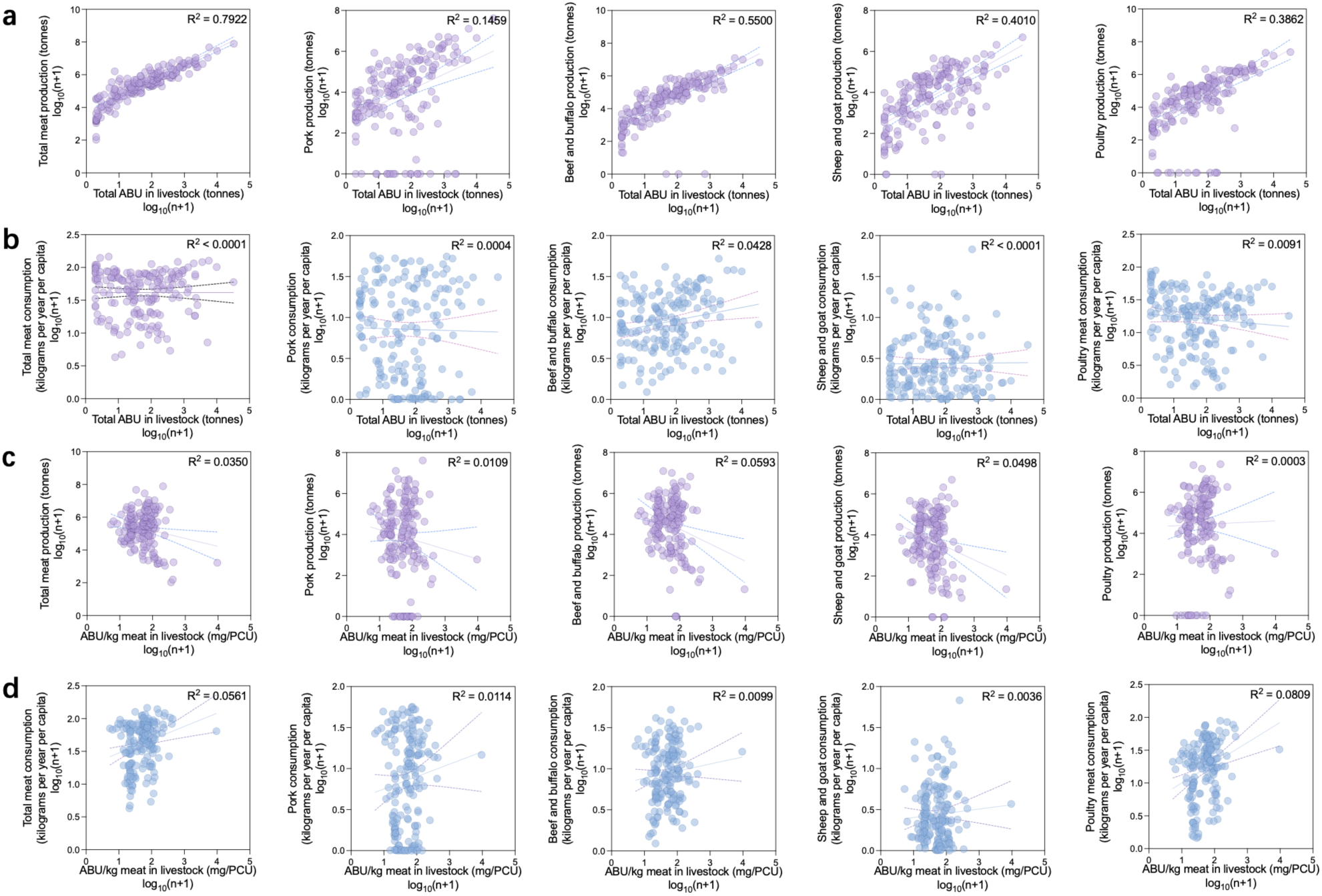
Associations between ABU in livestock and meat production or consumption. (a) Scatterplots showing the relationships between total ABU in livestock and production of total meat, pig meat, beef and buffalo meat, sheep and goat meat, and poultry meat. (b) Scatterplots showing the relationships between total ABU in livestock and consumption of total meat, pig meat, beef and buffalo meat, sheep and goat meat, and poultry meat. (c) Scatterplots showing the associations between ABU/kg meat in livestock and production of total meat and each meat type. (d) Scatterplots showing the associations between ABU/kg meat in livestock and meat consumption (total and by type). All variables were log ^(n+1)^-transformed for statistical analysis.

**Table 1.** Spearman correlation coefficients between ABU in livestock and meat production or consumption.

| Items | Total ABU in livestock |  | ABU/kg meat in livestock |  | N |
| --- | --- | --- | --- | --- | --- |
|  | r | p | r | p |  |
| Total meat production | **0.906** | **< 0.001** | 0.002 | 0.983 | 181 |
| Pork production | **0.493** | **< 0.001** | -0.055 | 0.461 | 181 |
| Beef and buffalo production | **0.849** | **< 0.001** | -0.083 | 0.264 | 181 |
| Sheep and goat production | **0.651** | **< 0.001** | -0.107 | 0.152 | 181 |
| Poultry production | **0.742** | **< 0.001** | 0.062 | 0.406 | 181 |
| Total meat consumption | -0.025 | 0.742 | **0.232** | **0.002** | 181 |
| Pork consumption | -0.031 | 0.674 | 0.09 | 0.23 | 181 |
| Beef and buffalo consumption | **0.199** | **0.007** | 0.092 | 0.216 | 181 |
| Sheep and goat consumption | 0.007 | 0.925 | 0.024 | 0.748 | 181 |
| Poultry consumption | -0.133 | 0.075 | **0.268** | **< 0.001** | 181 |

In contrast, total ABU in livestock was not associated with total meat consumption (r = -0.0825, *p* = 0.742), nor with the consumption of pig meat (r = -0.031, *p* = 0.674), sheep and goat meat (r = 0.007, *p* = 0.925), or poultry meat (r = -0.133, *p* = 0.075) (Figure 2b; Table 1).

Similarly, ABU intensity (ABU/kg meat in livestock) showed no significant correlation with total meat production (r = 0.002, *p* = 0.983), pig meat (r = -0.055, *p* = 0.461), beef and buffalo meat (r = -0.083, *p* = 0.264), sheep and goat meat (r = -0.107, *p* = 0.152), or poultry meat (r = 0.062, *p* = 0.406) (Figure 2c; Table 1). Nor was ABU/kg meat in livestock associated with consumption of pig meat (r = 0.090, *p* = 0.230), beef and buffalo meat (r = 0.092, *p* = 0.216) or sheep and goat meat (r = 0.024, *p* = 0.784) (Figure 2d; Table 1). However, it showed significant correlation with total meat consumption (r = 0.232, *p* = 0.002) and poultry meat consumption (r = 0.268, *p* < 0.001)

These findings indicate that increasing ABU is likely driven by production demands, as meat production scale appears tightly linked to antibiotic inputs. However, neither increased ABU nor increased meat production has been accompanied by improvements in ABU efficiency, as reflected by ABU/kg meat in livestock, nor have these increases been matched by consumption trends.

We further examined the relationships between ABU metrics and national economic and educational levels. Neither GDP per capita nor average LAYs were significantly associated with total ABU in livestock (GDP per capita: r = 0.097, *p* = 0.239; average LAYs: r = 0.157, *p* = 0.056). For ABU intensity (ABU/kg meat), GDP per capita remained non-significant (r = 0.110, *p* = 0.182), whereas average LAYs showed a weak but statistically significant positive correlation (r = 0.182, *p* = 0.027) (Figures S1a and S2a; Table S1). These results suggest that economic and educational advancement have not translated into more efficient ABU practices in livestock sectors. This could reflect insufficient or ineffective antimicrobial stewardship education for livestock producers, or limitations of general education in influencing sector-specific behaviors.

Interestingly, GDP per capita was significantly correlated with total meat consumption (r = 0.809, *p* < 0.001) and total meat production (r = 0.231, *p* = 0.005), as well as with consumption of pig meat (r = 0.564, *p* < 0.001), beef and buffalo meat (r = 0.583, *p* < 0.001), poultry meat (r = 0.676, *p* < 0.001) and pig meat production(r = 0.303, *p* < 0.001) (Figure S1b,d; Table S1). Education level (average LAYs) exhibited similar associations: positively correlated with total meat production (r = 0.286, *p* < 0.001), pig meat (r = 0.484, *p* < 0.001), beef and buffalo meat (r = 0.164, *p* = 0.046), and poultry meat (r = 0.172, *p* = 0.036) production, total meat consumption (r = 0.790, *p* < 0.001), pig meat (r = 0.714, *p* < 0.001), beef and buffalo meat (r = 0.551, *p* < 0.001), and poultry meat (r = 0.591, *p* < 0.001) consumption (Figure S2b,d; Table S1).

Moreover, given the strong correlation between GDP per capita and education level (r = 0.854, *p* < 0.001), the relationships between GDP per capita or education and meat production or consumption may be influenced by underlying collinearity between economic and developmental indicators. Overall, these results collectively suggest that while economic affluence and education are associated with higher consumption of animal-source foods, they do not necessarily translate into reduced antibiotic inputs or enhanced ABU efficiency in livestock production.

### 3.3 Global regulatory strategies on livestock ABU: heightened concern without enhanced efficiency

As established in Section 3.2, neither increases in total ABU nor growth in meat production or consumption were associated with improvements in ABU efficiency. While the total volume of ABU is inherently driven by the scale of livestock intensification and remains difficult to control, ABU intensity (ABU/kg meat) is, in theory, a more tractable and decoupled target. Thus, understanding whether current regulatory frameworks effectively improve ABU efficiency is of critical importance. In global antimicrobial policies, legislation often targets the broader category of antimicrobials rather than antibiotics specifically. Nonetheless, because antibiotics constitute the majority of antimicrobials used in animal agriculture, analyses based on antimicrobial regulation remain broadly informative.

As of 2020, among the 171 countries or regions with available data, 116 (67.84%) had legislation restricting antimicrobial use in livestock, while 55 (32.16%) had no such restrictions (Figure 3c). Interestingly, countries with antimicrobial use regulations reported significantly higher total ABU (*p* < 0.0001), as well as higher levels of meat production (*p* < 0.0001) and consumption (*p* < 0.01), compared to unregulated countries (Figure 3a). These regulated countries also had higher GDP per capita (*p* < 0.0001) and greater average years of schooling (*p* < 0.0001). This pattern suggests that regulation is more likely to be implemented in countries with high meat demand, intensive livestock systems, and greater public awareness of ABU risks.

**Figure 3.**
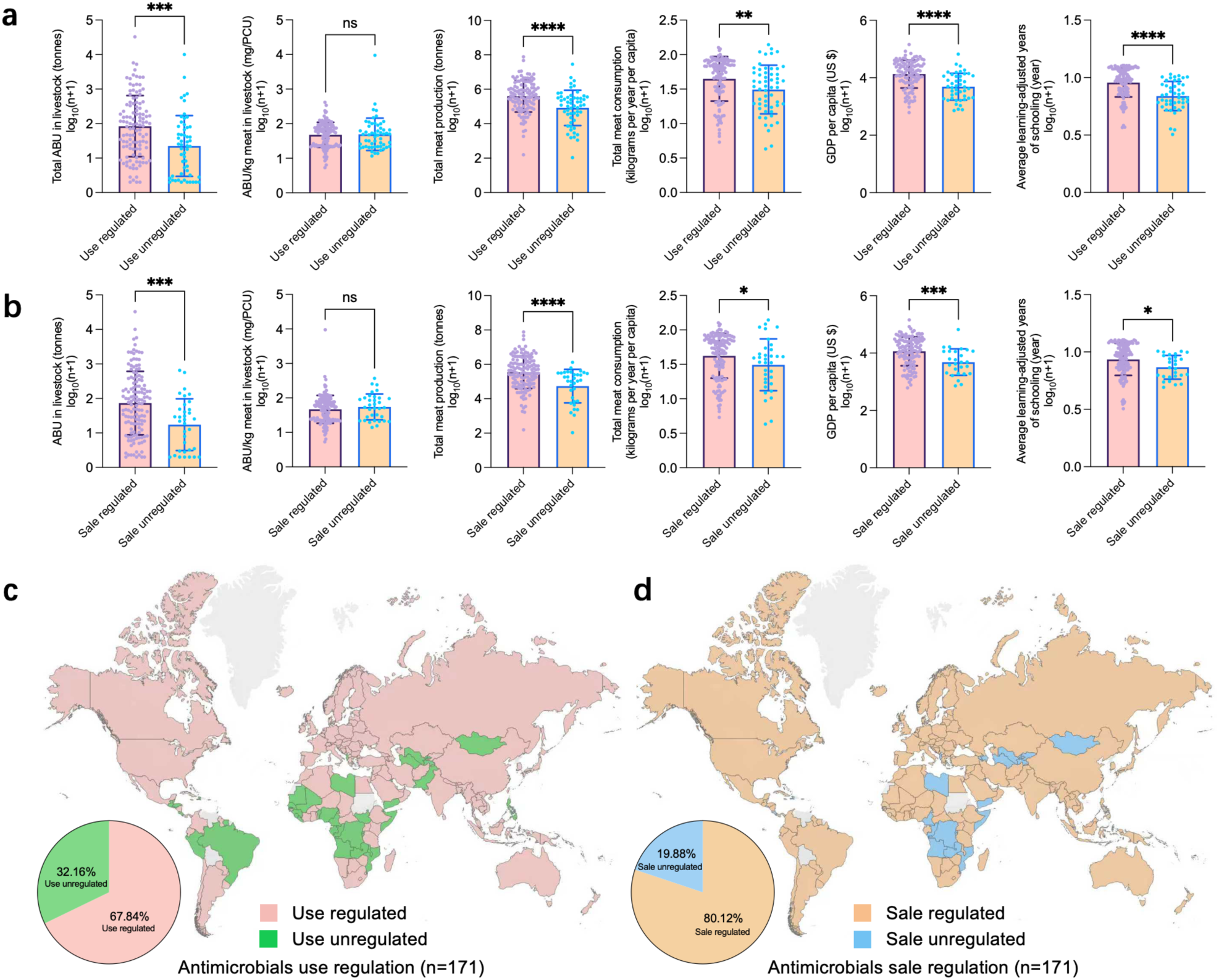
Global regulatory landscape of antimicrobial use and antimicrobial sales in livestock production. (a) Comparisons of total ABU in livestock, ABU/kg meat in livestock, total meat production and consumption, GDP per capita, and average LAYs between countries or regions with and without legislation restricting antimicrobial use in livestock. (b) Comparisons of the same indicators between countries or regions with and without legal frameworks regulating the sale of antimicrobials for livestock. (c) Global distribution of countries or regions with legal prohibitions on the antimicrobials use for growth promotion in livestock in the absence of prior risk analysis. (d) Global distribution of countries or regions with laws or regulations governing the sale of antimicrobials for terrestrial animal production.

However, there was no significant difference in ABU efficiency (ABU/kg meat) between regulated and unregulated countries. This surprising observation implies that despite a stronger regulatory presence in high-use settings, these measures have not translated into greater efficiency. This could reflect limited enforcement, superficial compliance, or the absence of targeted interventions focused specifically on efficiency metrics.

A similar pattern was observed for legislation governing the sale of antimicrobials. Among the same 171 countries, 137 (80.12%) had legal frameworks regulating antimicrobial sales for livestock, while 34 (19.88%) did not (Figure 3d). Although such sales regulation should, in principle, exert more direct control over access to antibiotics and thereby potentially reduce overuse, countries implementing these regulations nevertheless exhibited significantly higher meat production (*p* < 0.0001), consumption (*p* < 0.05), total ABU (*p* < 0.001), GDP per capita (*p* < 0.0001), and education levels (*p* < 0.05) compared to those without such policies (Figure 3b).

These findings suggest that both regulatory approaches, including usage restrictions and sales controls, are more commonly adopted in countries with higher levels of meat consumption, GDP per capita, and educational attainment. This pattern may reflect a greater societal awareness of antibiotic resistance or a stronger sense of social responsibility in these regions. However, these policies have not yet produced measurable gains in efficiency, and no significant differences were observed between the two regulatory approaches, despite the theoretical advantages of sales-based control.

In summary, global antimicrobial regulations appear to be reactive to concerns over escalating ABU, particularly in high-output regions. However, their impact remains limited, suggesting that existing frameworks may lack the specificity, enforcement, or integration with behavioral and technical interventions necessary to drive meaningful improvements in antibiotic stewardship.

### 3.4 Regional patterns and trends in ABU

#### 3.4.1 Continental variation in factors associated with ABU

While previous analyses (Sections 3.1 and 3.2) elucidated global trends in ABU in livestock, regional heterogeneity necessitates continent-specific examinations to uncover localized factors influencing ABU dynamics. Given cultural similarities, comparable economic statuses, geographic proximities, and common livestock management practices within continents, we conducted detailed intra-continental analyses for Asia, Europe, North America, South America, Africa, and Oceania.

Intriguingly, continent-level patterns differed markedly from global observations (Figure 4, S3-S5; Table S2-S4). In Asia and Europe, total ABU in livestock demonstrated significant positive correlations with ABU intensity (ABU/kg meat), indicating that higher absolute usage coincided with lower antibiotic efficiency. Conversely, North America exhibited a significant negative correlation, suggesting improved antibiotic efficiency concurrent with higher absolute usage. In South America, Africa, and Oceania, no significant relationship between total ABU and ABU intensity emerged. These findings highlight substantial geographical disparities in antibiotic stewardship, emphasizing that in regions with intensive antibiotic practices, particularly Asia and Europe, high usage does not equate to improved efficiency.

**Figure 4.**
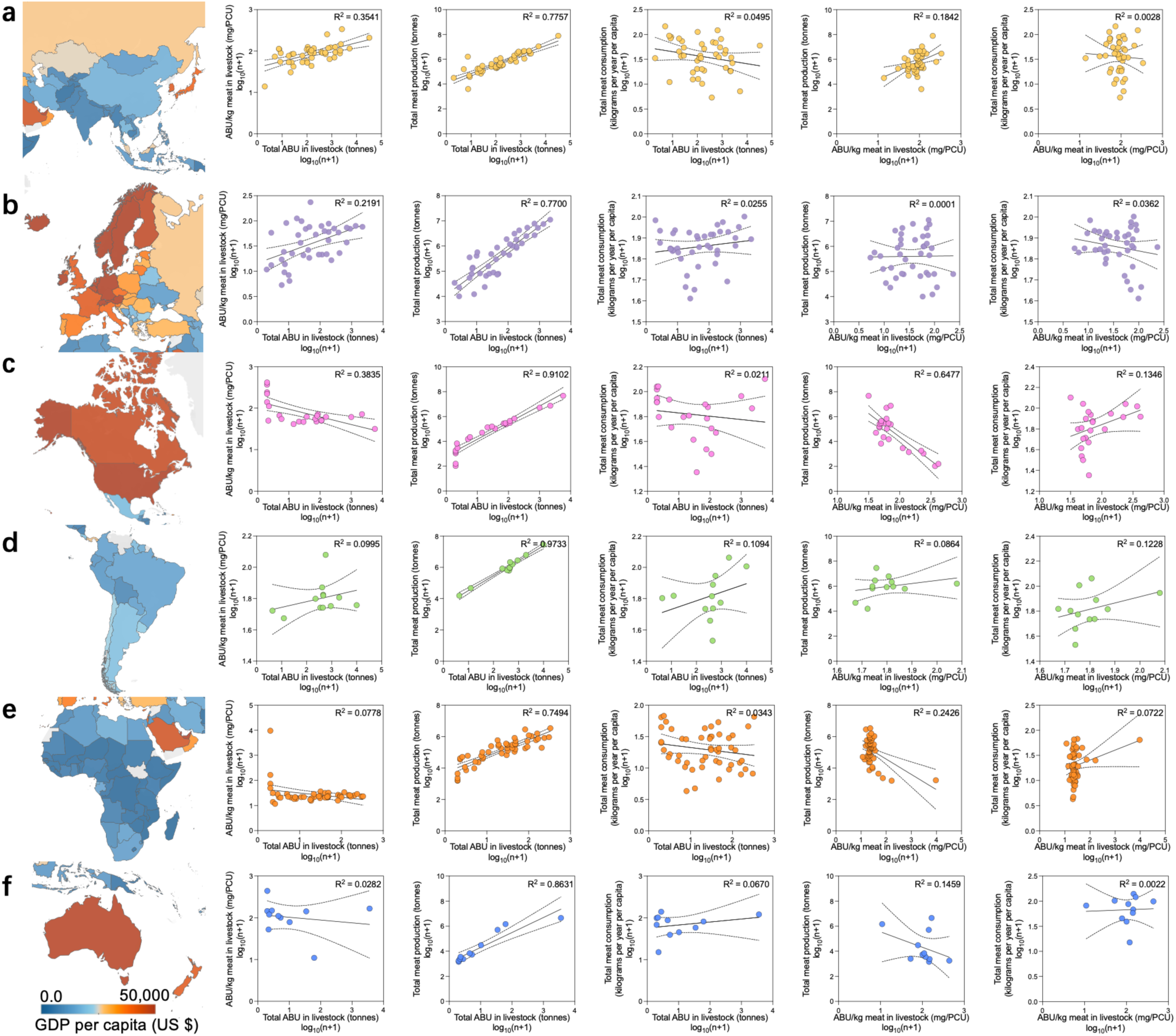
Within-continent associations between ABU in livestock and meat production and consumption. Panels a-f correspond to Asia, Europe, North America, South America, Africa, and Oceania, respectively.

Consistent with global trends, total ABU in livestock across all continents strongly correlated with total meat production, reaffirming the universal dependency of meat production systems on antibiotic inputs. This persistent reliance across diverse regional contexts suggests meat production growth fundamentally drives antibiotic demand, rather than solely reflecting inefficiencies in antibiotic stewardship.

Notably, ABU intensity exhibited negative correlations with meat production in North America and Africa. In particular, the North American trend, characterized by rising antibiotic efficiency alongside increased production, underscores potential successes in antibiotic stewardship that could inform global practices.

No significant differences emerged in other analyzed parameters between continental and global scales, underscoring consistency in the broader relationships among antibiotic usage, meat production, and consumption patterns.

#### 3.4.2 Temporal changes in antibiotic efficiency and meat production in Europe

European countries, characterized by advanced livestock management practices, stringent antibiotic regulatory frameworks, and high standards of production, present an ideal context for evaluating changes in antibiotic efficiency. Given the region’s robust data infrastructure and detailed historical records, we analyzed temporal trends in antibiotic efficiency from 2010 to 2022 across 31 European countries, alongside concurrent meat production, consumption, and GDP per capita trends from 2000 to 2022 (Figure 5).

**Figure 5.**
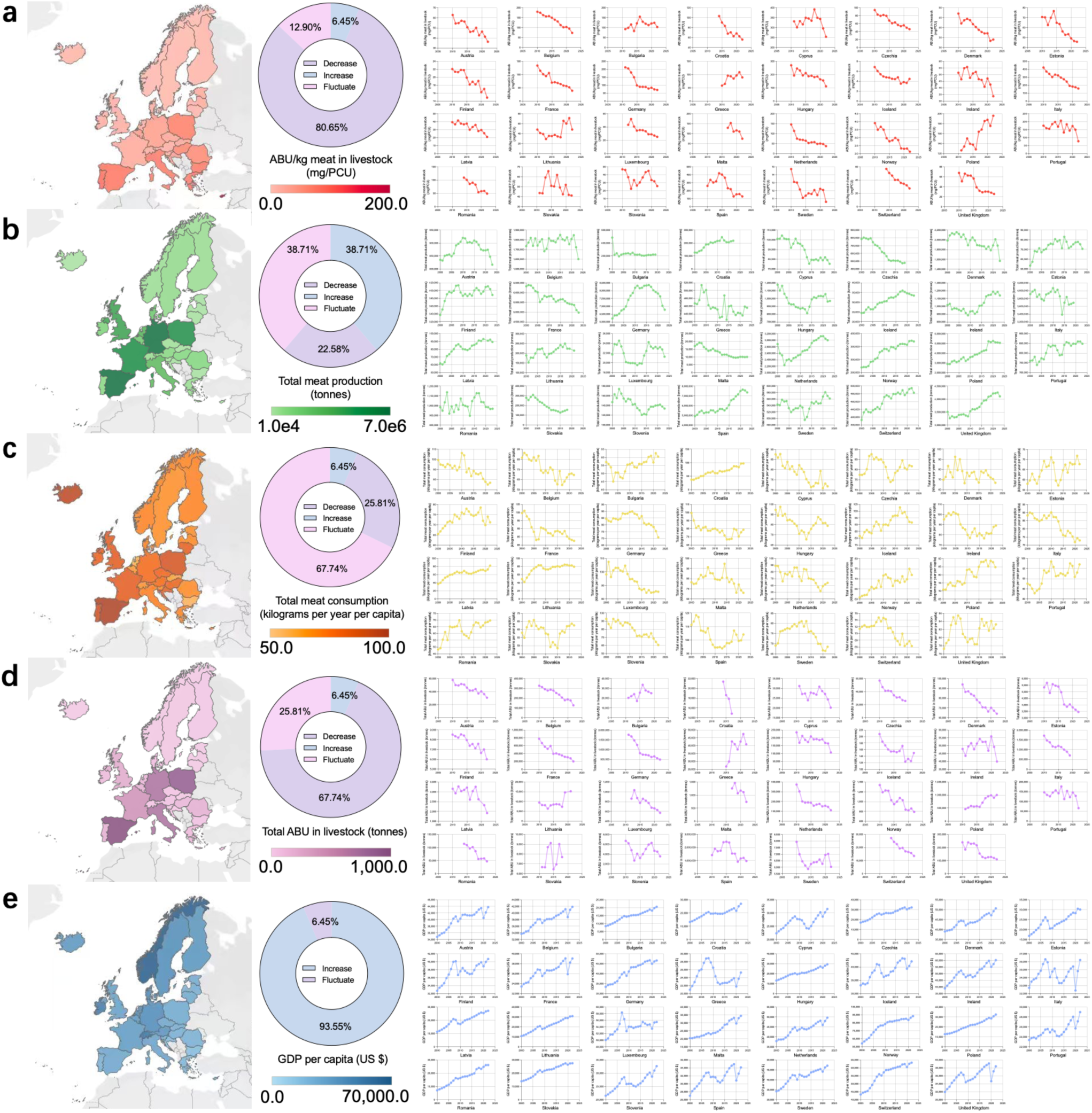
Temporal trends in ABU and meat production and consumption in Europe. (a) ABU/kg meat in livestock in livestock across European countries from 2010 to 2022. (b) Changes in total meat production from 2000 to 2022. (c) Changes in total meat consumption from 2000 to 2022. (d) Changes in GDP per capita from 2000 to 2022. (e) Estimated total ABU in livestock in Europe from 2010 to 2022.

Strikingly, 80.65% of analyzed countries exhibited declining trends in ABU intensity (ABU/kg meat), indicating widespread improvements in antibiotic efficiency (Figure 5a). A minority of countries demonstrated fluctuating (12.90%) or increasing (6.45%) trends. Correspondingly, total ABU declined significantly in 67.74% of countries, with 25.81% showing fluctuations and only 6.45% indicating increases (Figure 5d). Crucially, these reductions in total ABU occurred without parallel declines in total meat production (Figure 5b). Indeed, several countries, notably Iceland, Norway, and Switzerland, experienced significant production increases.

These findings underscore Europe’s potential to decouple meat production growth from antibiotic usage through enhanced efficiency and stringent stewardship practices, even amid increasing economic affluence (Figure 5c, e). This successful decoupling highlights viable pathways for global adoption, demonstrating that effective antibiotic governance need not constrain livestock productivity.

### 3.5 Global projections of ABU in livestock agriculture

Despite considerable global efforts to monitor ABU in food animal production, the historical trajectory and long-term projection of total livestock-related ABU remain poorly characterized, primarily due to incomplete national surveillance and inconsistent data availability across countries. To address this gap, we developed a robust forecasting framework leveraging a key empirical observation: the strong, consistent linear association between total meat production and ABU across countries. This relationship allows meat production to serve as a proxy indicator for ABU in data-deficient regions, providing a scalable approach for historical reconstruction and future prediction.

Using data from 1961 to 2023, we first modeled global total meat production over time. The linear regression model is expressed as:

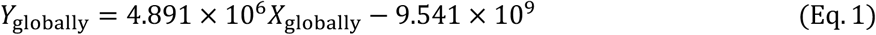

indicating that global meat output has increased at an average rate of approximately 4.89 million tonnes per year. We next applied a previously derived regression linking ABU and meat production in 2020:

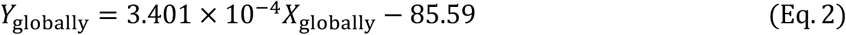

where each tonne of meat corresponds to 340.1 mg of antibiotics per population correction unit (mg/PCU). By combining Equation (Eq.) 1 and Eq. 2, we derived a global ABU prediction model as a function of time:

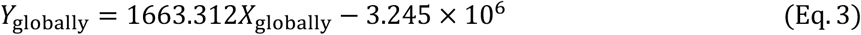

This indicates an alarming trend: global antibiotic consumption in livestock is increasing linearly at approximately 1,663 tonnes per year. Historical backcasting suggests that ABU began rising in the early 1950s, aligning with the introduction of antibiotics into agriculture around 1952. Our model estimates total livestock ABU reached 81,929 tonnes in 2000, 98,562 tonnes in 2010, and 115,195 tonnes in 2020. The increase from 2000 to 2020 alone amounts to over 33,000 tonnes. Future projections suggest total consumption may reach 131,828 tonnes in 2030 and 165,094 tonnes by 2050.

Cumulatively, we estimate that between 1950 and 2020, livestock production consumed approximately 4.05 million tonnes of antibiotics. Alarmingly, the period from 2020 to 2050 alone is projected to add a further 4.23 million tonnes, which exceeds the cumulative usage over the entire previous 70 years. By 2050, cumulative agricultural ABU could surpass 8.28 million tonnes globally. These trends underscore an urgent need for regulatory and technological interventions to curb excessive ABU in the livestock sector (Figure 6a).

**Figure 6.**
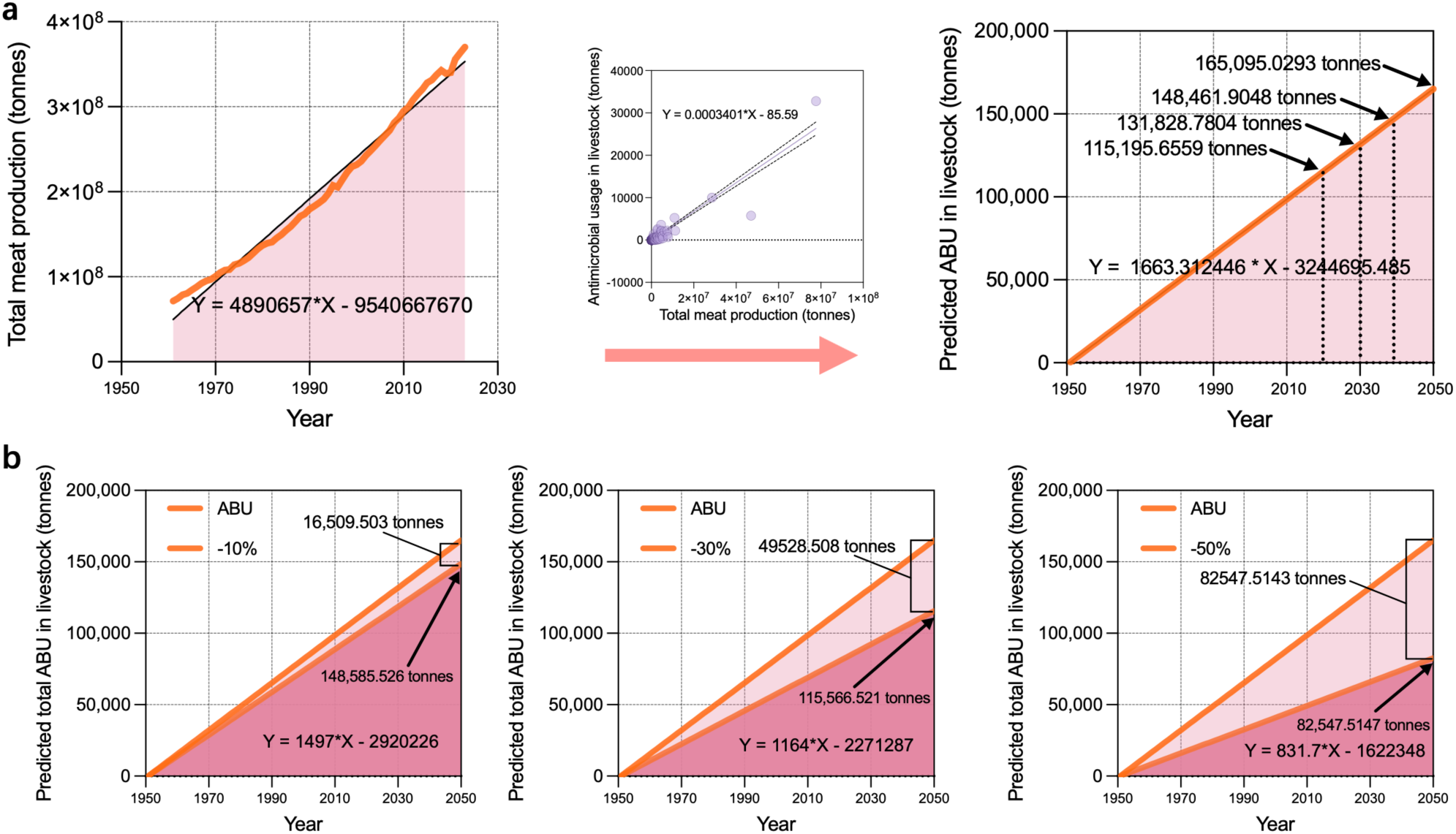
Global trends and future projections of total ABU in livestock, 1950-2050. (a) Projected global total ABU in livestock based on historical and predicted trends in total meat production and ABU efficiency (ABU/kg meat). (b) Projected global trajectories of total ABU in livestock from 1950 to 2050 under scenarios where annual growth rates in total ABU are reduced by 10%, 30%, or 50%.

To refine our projections, we also analyzed five major livestock-producing nations, namely China, the United States, Brazil, Canada, and Australia, which are characterized by high ABU levels and nearly linear growth in meat production. Their respective regression models for meat production over time are:

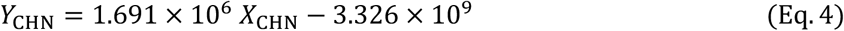

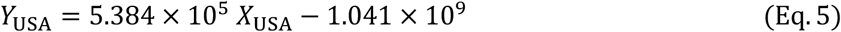

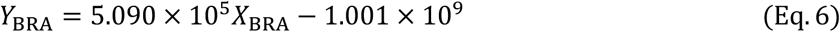

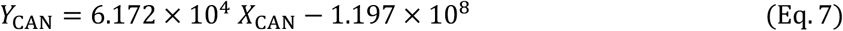

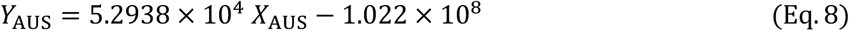

Integrating these with Eq. 2, we derived each country’s ABU forecast function:

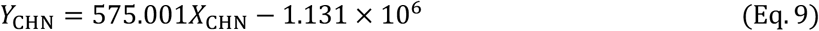

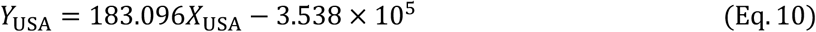

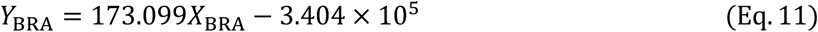

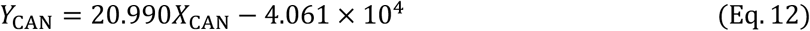

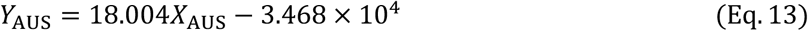

Accordingly, by 2025, projected livestock ABU is expected to reach 47,672 tonnes in China, 21,555 tonnes in the USA, 14,473 tonnes in Brazil, 2,416 tonnes in Canada, and 2,233 tonnes in Australia (Figure S7a, b).

Although cross-country heterogeneity in ABU efficiency is substantial, these differences have not yet disrupted the global ABU-meat production correlation. As such, meat output remains a useful surrogate for predicting ABU, particularly for countries lacking direct surveillance. This finding holds important implications for global antibiotic governance and the estimation of unreported consumption.

To explore mitigation potential, we modeled scenarios of improved ABU efficiency at global and national scales. Assuming annual ABU growth rates are reduced by 10%, 30%, or 50%, we estimated that by 2050, global ABU could decrease by 16,509 tonnes, 49,529 tonnes, and 82,548 tonnes, respectively (Figure 6b). For China, a 10% efficiency improvement would reduce ABU by ∼3,617 tonnes in 2030 and 4,767 tonnes in 2050; in the USA, the corresponding reductions would be ∼1,789 and 2,156 tonnes. Over the 2030–2050 period, this would result in cumulative ABU reductions of 88,036 tonnes in China and 41,421 tonnes in the United States, with each figure exceeding the total global ABU recorded in 2020.

Finally, given the strong and consistent ABU-meat production relationship (Eq. 2), this model offers a valuable framework for inferring spatiotemporal trends in ABU based on meat production data, which are far more widely available. For instance, time-series data from OWID provide detailed country-level records of meat production from 1961 onward, enabling retrospective mapping of antibiotic trends even in the absence of direct ABU statistics.

## 4. Discussion

Meeting surging demand for animal-sourced food has driven the rapid intensification of livestock production worldwide^19^. High-density systems reduce unit costs and stabilize supply, but they also heighten susceptibility to infectious disease, making antibiotics a routine input for prophylaxis and metaphylaxis^20,21^. The resulting scale of ABU in livestock is substantial and, more importantly, often misaligned with principles of evidence-based stewardship. Our study provides a comprehensive evaluation of the global ABU landscape in livestock systems. By integrating national meat production as a pragmatic proxy, we systematically quantified historical patterns, dissected the socioeconomic and regulatory drivers of use intensity, and revealed substantial spatial heterogeneity in usage efficiency^22,23^. This integrative approach bridges critical data gaps, informs policy relevance, and underpins subsequent projections to 2050. Together, these analyses illuminate both the structural determinants of antibiotic overuse and actionable opportunities for mitigation within a One Health framework^24–26^.

### 4.1 Global landscape and efficiency gaps of ABU in livestock

We document pronounced cross-country heterogeneity: a small number of nations account for a large share of global ABU, while ABU intensity varies by orders of magnitude. Countries with the greatest total ABU tend to exhibit lower efficiency, indicating persistent reliance on antibiotics to sustain production rather than on biosecurity, vaccination, precision nutrition, or husbandry improvements^16,27^. At the continental scale, patterns diverge: Asia and Europe demonstrate a positive association between total ABU and usage intensity; North America exhibits a negative association that reflects improved efficiency; Africa, South America, and Oceania show no consistent pattern, which may be attributed to heterogeneous production systems and limitations in data availability. Notably, within Europe (2010–2022), 80.65% of countries reduced ABU intensity and 67.74% reduced total ABU while maintaining or increasing output, demonstrating that efficiency gains are achievable without sacrificing production.

### 4.2 Global drivers of inefficient ABU in livestock

A striking insight from our analysis is that higher national wealth or education levels do not automatically translate into more efficient ABU. We found no significant negative association between ABU efficiency and socioeconomic indicators such as GDP per capita or average educational attainment^16,28^. This counterintuitive result suggests that development and awareness alone are insufficient to ensure prudent antibiotic practices in livestock. Instead, inefficiencies in ABU appear to be driven by structural and economic factors inherent to modern animal production systems^16,29^. Intensification of livestock farming is a key underlying driver: high-density animal housing creates constant disease pressures, incentivizing routine prophylactic ABU as a low-cost insurance policy against outbreaks^5,30^. In many cases, it is economically more convenient for producers to rely on antibiotics, which are often inexpensive and effective in the short term, rather than to invest in improved biosecurity, sanitation, or vaccination programs^31,32^. Widespread availability of over-the-counter veterinary drugs and insufficient farmer training in best practices further exacerbate indiscriminate use of antimicrobials in numerous regions^33^. Even as public concern about food safety and antibiotic residues has grown in some countries, this has not consistently translated into better on-farm practices^34^. In other words, there remains a considerable gap between recognizing the importance of antibiotic stewardship and implementing the husbandry changes required to achieve it^35^. These global drivers of inefficiency ensure that, absent strong interventions, the misuse of antibiotics in livestock can persist even in settings where awareness is high^11,29^.

### 4.3 Policy-implementation gaps and regulatory limitations

Our analysis reveals a critical disconnect between formal regulatory presence and actual improvements in ABU efficiency. Although countries implementing use-side or sales-side regulatory frameworks often have higher meat production, stronger economies, and better educational levels, these policies alone are not linked to more prudent ABU^36,37^.

Several systemic barriers likely underpin this ineffectiveness. Use-side regulations require costly enforcement infrastructures such as veterinary oversight, on-farm inspections, and routine residue testing, which are inconsistently implemented and suffer from fragmented national standards^17,38,39^. Sales-side controls, while potentially limiting upstream availability, are vulnerable to circumvention through informal markets, online vendors, or regulatory loopholes, and often lack real-time enforcement mechanisms^40,41^. Moreover, both approaches tend to be reactive rather than preventive, and rely on legal instruments whose effects are lagged and indirect^17,37^.

These limitations underscore that legislation, though necessary, is insufficient in isolation. Effective governance must operate across multiple levels, combining enforceable regulatory standards with proactive system enablers: market-based incentives, professional education, real-time surveillance systems, and cost-effective technologies that reduce the burden of compliance^16,28,42^. Ultimately, successful stewardship depends not only on rule-making, but on reducing the systemic friction of “doing the right thing” within agrifood systems.

### 4.4 Projected trajectories to 2050 and the mitigation potential of efficiency

Our projection exercise provides a methodological extension to previous research by anchoring forecasts to globally available meat production data and an empirically derived coefficient linking production to ABU^16,43,44^. This approach enables the reconstruction of livestock antibiotic consumption from the early 1950s and extends projections to 2050, a temporal horizon rarely explored by earlier short-term studies^16,42^. This unified proxy framework yields a coherent global picture and reveals a nearly linear increase in ABU in livestock, with an average annual growth of approximately 1,663 tonnes. By 2050, the projected total is expected to reach around 165,000 tonnes, a level that aligns with earlier forecasts while encompassing a broader temporal span (Figure 6). The cumulative burden is striking: projected use over the next three decades surpasses that of the previous seven, underscoring the systemic nature of antibiotic dependence in modern livestock systems.

Interpreted through a policy lens, the projections locate where leverage lies. Growth in a handful of major producers (e.g., China, USA, Brazil, Canada, Australia) accounts for much of the future increase (Figure S7), so targeted action can bend the global curve disproportionately. Critically, efficiency-led mitigation scales with production: reducing the annual growth rate of total ABU by 10%, 30%, or 50% by 2050 would result in a global ABU reduction of approximately 16,500 to 82,500 tonnes in that year, highlighting how moderate but widespread efficiency gains can yield substantial absolute decreases in antibiotic consumption (Figure 6).

While some simplifications are inherent to our projection strategy, they reflect the considerable global variability in ABU efficiency, species composition, production systems, and veterinary access, particularly in countries where routine ABU reporting is absent^29,43^. These factors constrain point-level precision and caution against overinterpretation at the national scale. The use of a fixed 2020 coefficient and a linear extrapolation model may also introduce limitations. Nevertheless, this approach addresses a critical gap in surveillance by leveraging a transparent and empirically derived linkage between production volume and ABU. It establishes a reproducible reference framework that can be locally adapted, encouraging countries to initiate ABU tracking and report both absolute quantities and usage intensity over time^16,29,43^. Importantly, this work underscores the feasibility of establishing a standardized global accounting system for ABU in livestock and offers a practical foundation for future improvements, including species-level resolution and policy-impact evaluations^45–47^.

### 4.5 From evidence to action: an integrated roadmap

Building on the observed production-ABU relationship, we propose an integrated strategy to control livestock ABU through upstream source control and downstream governance (Figure 7). Upstream, control measures should restrict unnecessary antibiotic availability and accelerate the deployment of validated alternatives^36,48^. These include regulated licensing with production caps, withdrawal of over-the-counter sales, fiscal alignment with stewardship goals, and promotion of substitutes such as vaccines, immunomodulators, probiotics or synbiotics, bacteriophage therapy, and precision feed additives^49–51^. Downstream, measures must ensure responsible on-farm and market practices through enhanced prescription oversight, digital audit trails supported by rapid diagnostics, harmonized biosecurity and animal welfare standards, routine antibiotic residue surveillance, and transparent product labeling that enables procurement decisions favoring low-ABU systems^52–56^.

**Figure 7.**
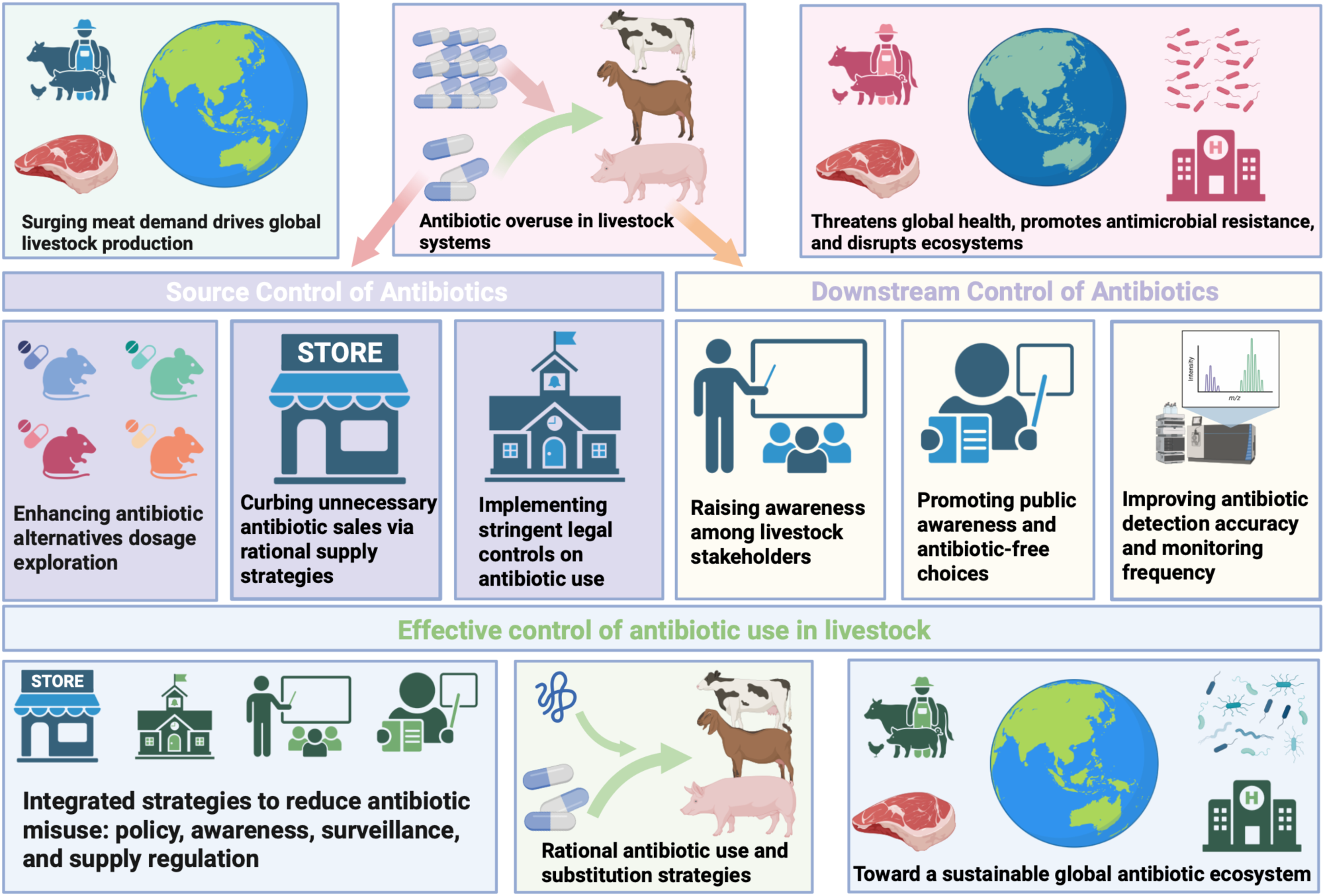
Source control and downstream governance strategies for livestock ABU.

As foundational system enablers, governments should institutionalize regular ABU monitoring using meat production as a proxy where direct measurements are lacking, publish open-access dashboards reporting both total and intensity metrics, and invest in the training of veterinarians and producers. These policy tools should be paired with performance-based incentives to accelerate uptake^46,57,58^. When integrated across the supply chain, this evidence-driven roadmap offers a credible and scalable pathway for reducing global antibiotic dependence and flattening the ABU trajectory well before mid-century (Figure 6 and 7).

## RESOURCE AVAILABILITY

### Lead contact

Further information and reasonable requests for resources and reagents should be directed to and will be fulfilled by the lead contact, Hongyue Ma.

### Materials availability

This study did not generate new samples or unique reagents.

#### Data and code availability

- ***Data***: This paper analyzes existing, publicly available data, accessible at [Database: https://ourworldindata.org].
- ***Code***: This paper does not report original code.
- ***Additional Information***: Any additional information required to reanalyze the data reported in this article is available from the lead contact upon request.

## ACKNOWLEDGMENTS

None.

## AUTHOR CONTRIBUTIONS

Y.L. and H.M. contributed equally to this work. They performed all analyses and wrote the manuscript.

## Supplementary Materials

**Figure S1.**
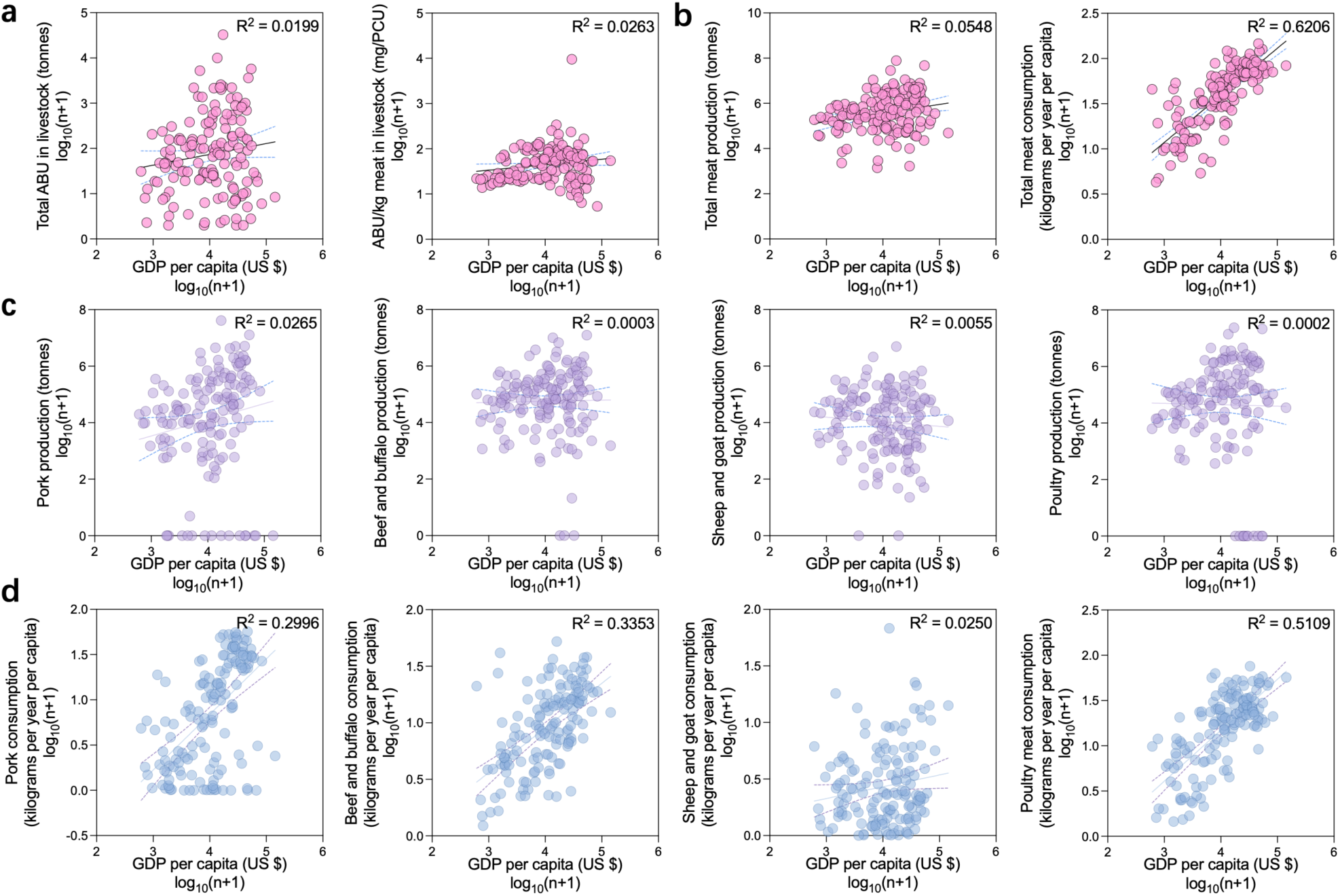
Associations between GDP per capita and ABU in livestock and meat production or consumption. (a) Scatterplots showing the associations between GDP per capita and both total ABU in livestock and ABU/kg meat in livestock. (b) Scatterplots showing the associations between GDP per capita and total meat production and consumption. (c) Scatterplots showing the associations between GDP per capita and production of pig meat, beef and buffalo meat, sheep and goat meat, and poultry meat. (d) Scatterplots showing the associations between GDP per capita and consumption of pig meat, beef and buffalo meat, sheep and goat meat, and poultry meat. All variables were log_10_^(n+1)^-transformed for statistical analysis.

**Figure S2.**
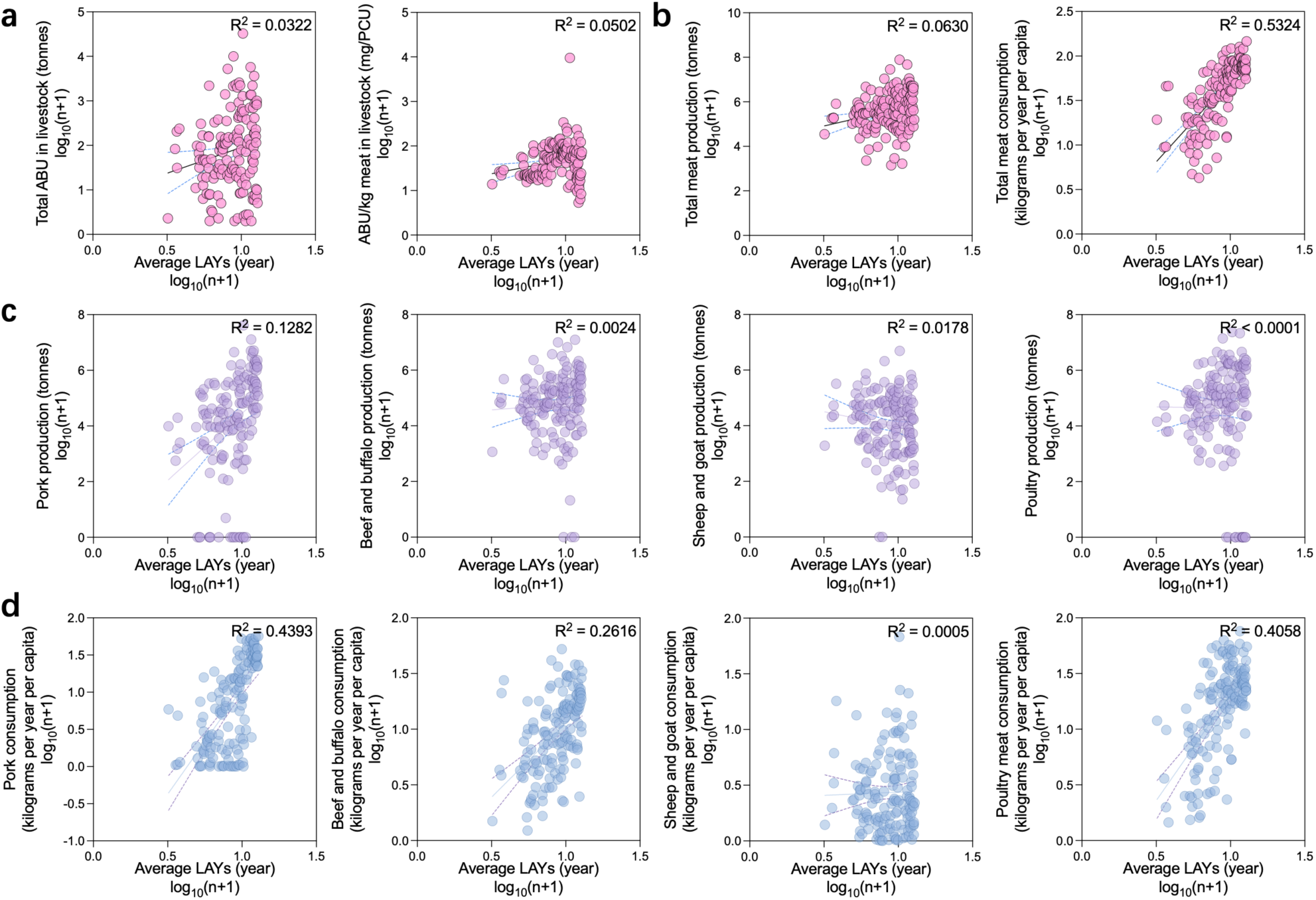
Associations between average LAYs and ABU in livestock and meat production or consumption. (a) Scatterplots showing the associations between average LAYs and both total ABU in livestock and ABU/kg meat in livestock. (b) Scatterplots showing the associations between average LAYs and total meat production and consumption. (c) Scatterplots showing the associations between average LAYs and production of pig meat, beef and buffalo meat, sheep and goat meat, and poultry meat. (d) Scatterplots showing the associations between average LAYs and consumption of pig meat, beef and buffalo meat, sheep and goat meat, and poultry meat. All variables were log_10_^(n+1)^-transformed for statistical analysis.

**Figure S3.**
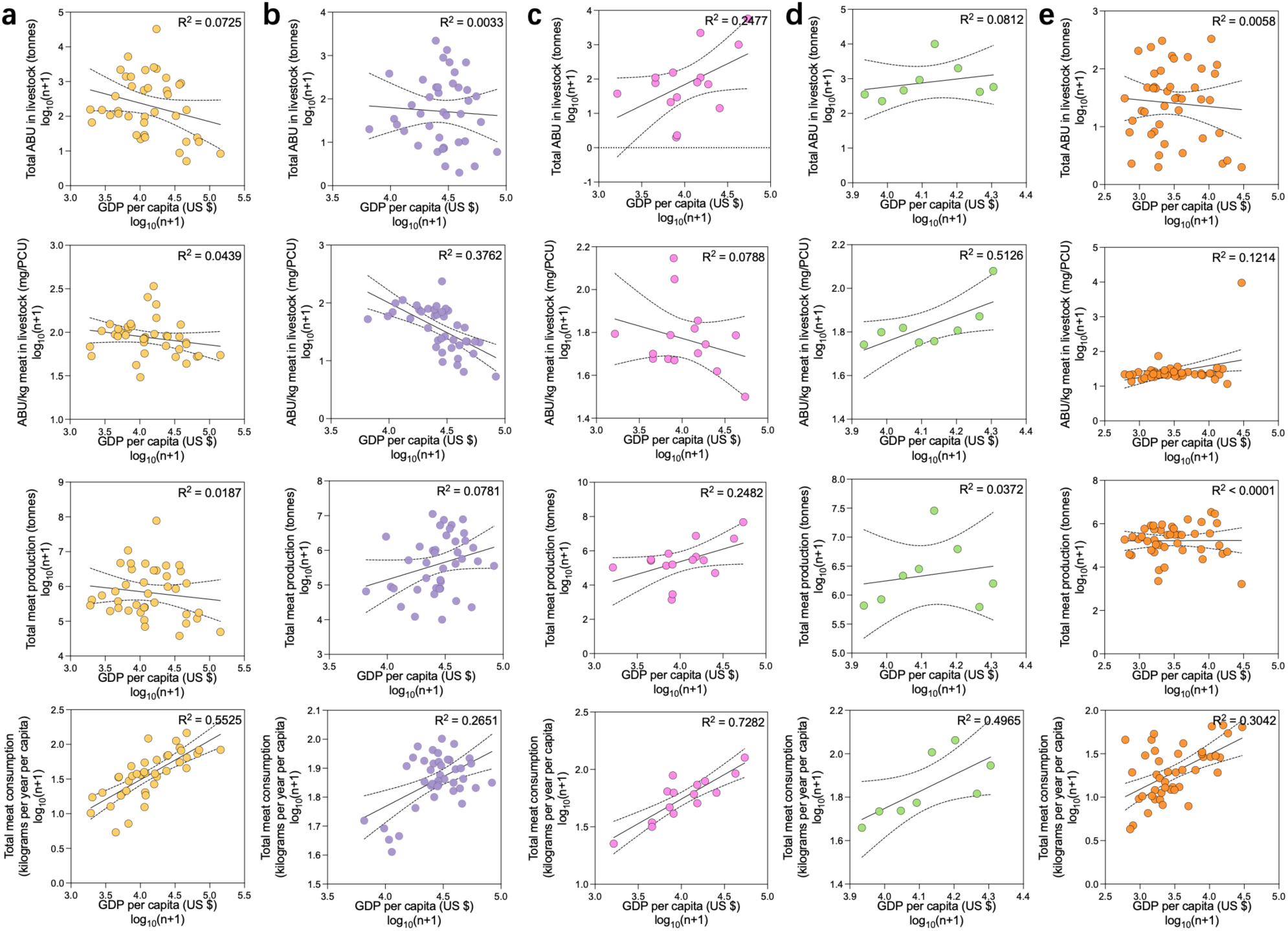
Associations between GDP per capita and ABU in livestock, total meat production or consumption across five continents. Panels a-e correspond to Asia, Europe, North America, South America and Africa, respectively. Due to insufficient data, Oceania was not included in the calculations. All variables were log ^(n+1)^-transformed for statistical analysis.

**Figure S4.**
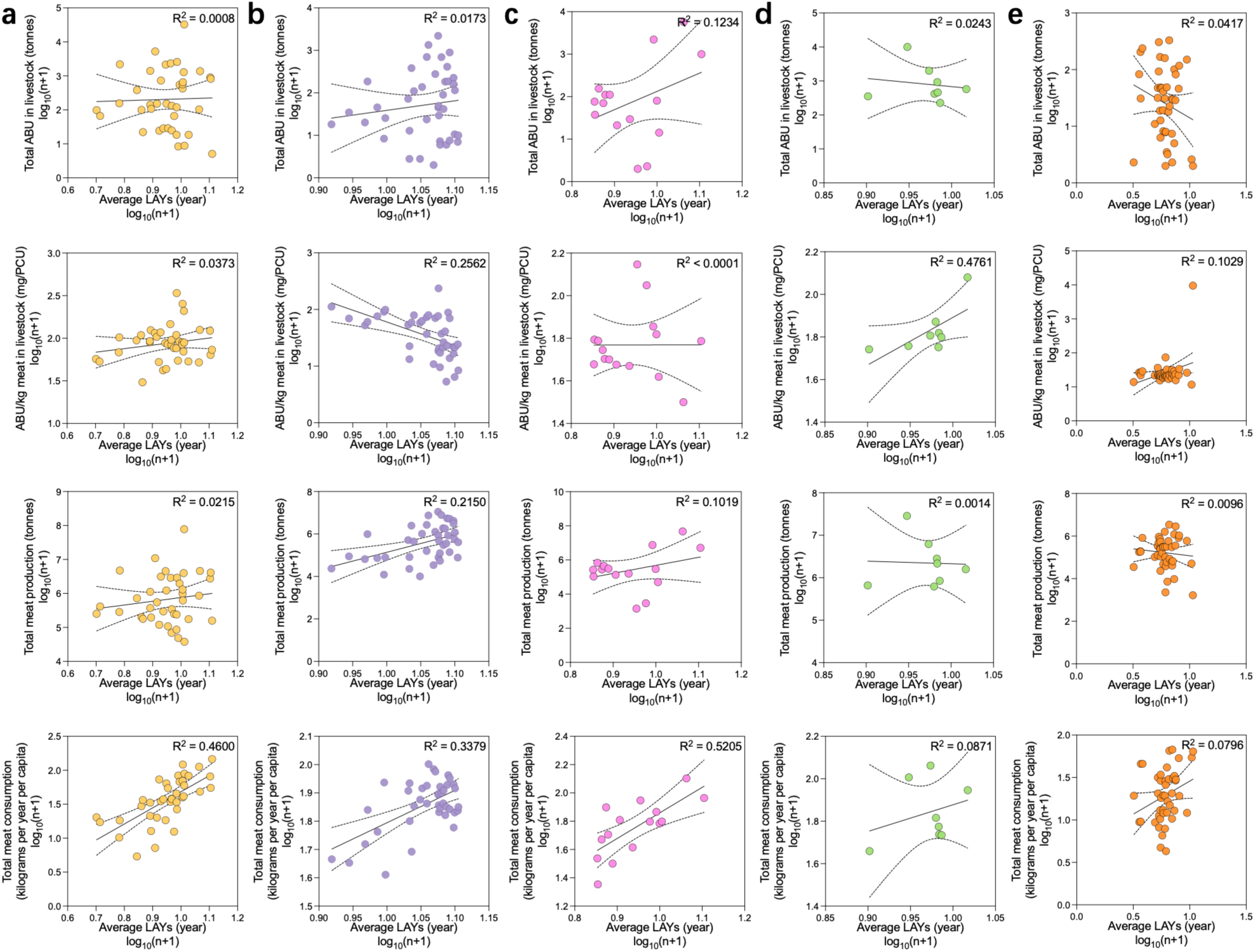
Associations between average LAYs and ABU in livestock, total meat production or consumption across five continents. Panels a-e correspond to Asia, Europe, North America, South America and Africa, respectively. Due to insufficient data, Oceania was not included in the calculations. All variables were log ^(n+1)^-transformed for statistical analysis.

**Figure S5.**
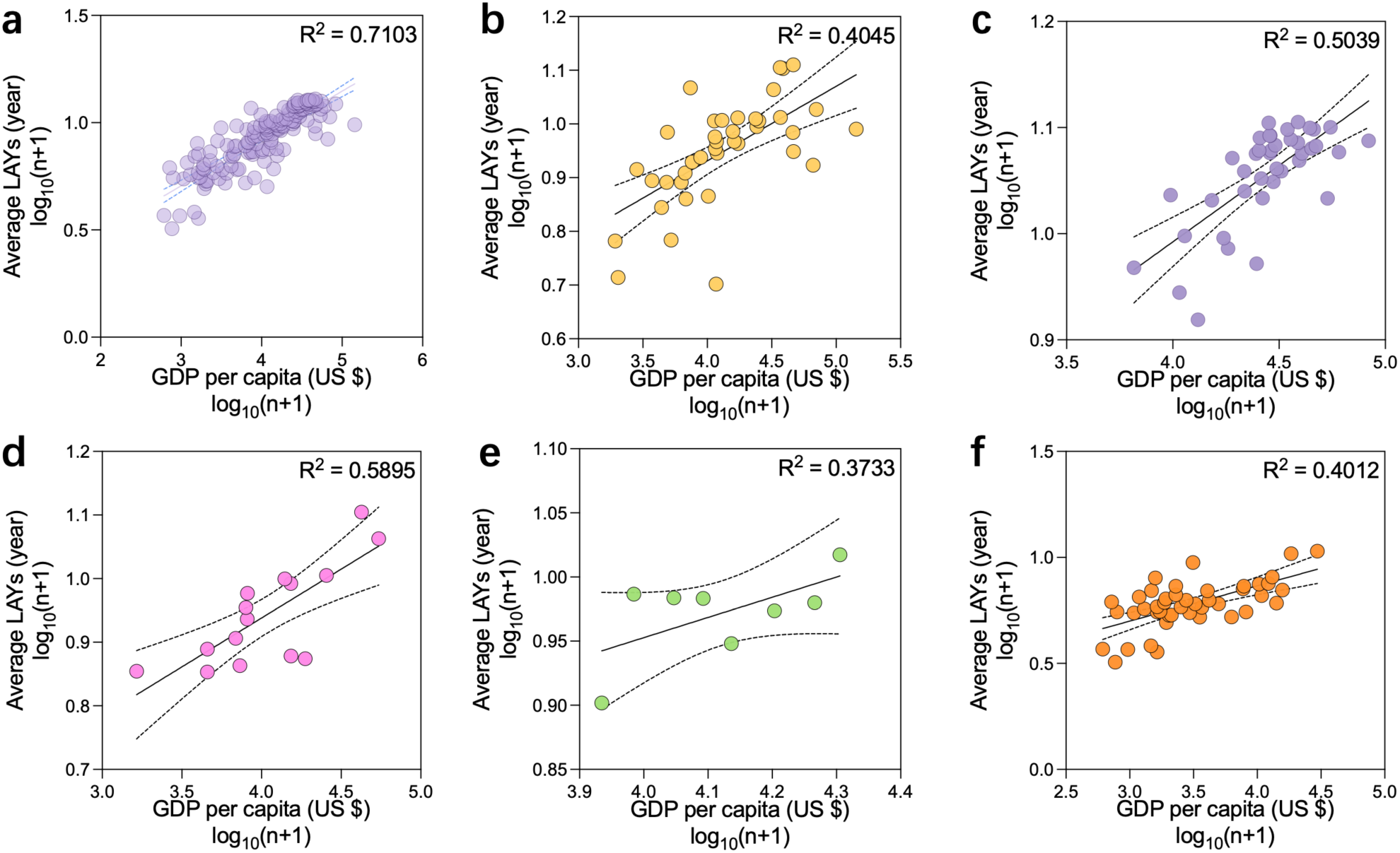
Associations between GDP per capita and average LAYs across global and continental scales. Panels a-f show the relationships for the global scale, Asia, Europe, North America, South America, and Africa, respectively. Due to insufficient data, Oceania was not included in the calculations. All variables were log_10_^(n+1)^-transformed for statistical analysis.

**Figure S7.**
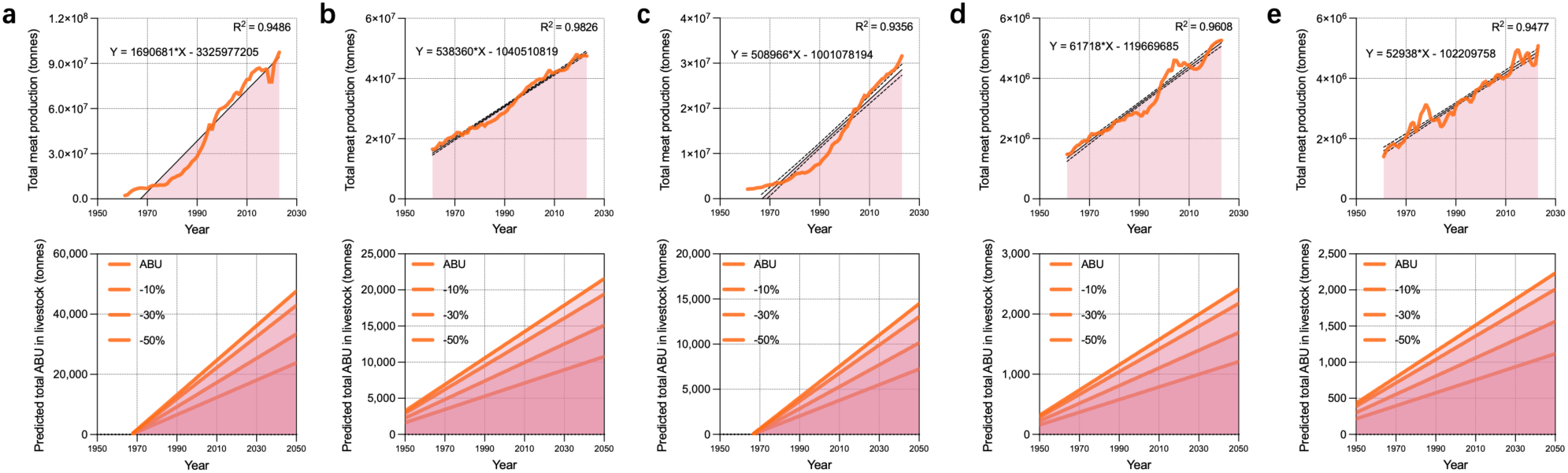
Projected trends and mitigation scenarios of ABU in livestock across representative countries or regions from 1960 to 2050. (a) Modeled projections of total meat production. (b) Estimated trajectories of total ABU in livestock under scenarios where annual total ABU growth rates are reduced by 10%, 30%, or 50% relative to current trajectories. Panels a-f correspond to China, the United States, Brazil, Canada, and Australia, respectively.

**Table S1.** Spearman correlation coefficients between GDP per capita or average LAYs and ABU in livestock, as well as total meat production or consumption.

| Items | Average LAYs (years) |  | GDP per capita |  | N |
| --- | --- | --- | --- | --- | --- |
|  | r | p | r | p |  |
| Total ABU in livestock | 0.157 | 0.056 | 0.097 | 0.239 | 148 |
| ABU/kg meat in livestock | **0.182** | **0.027** | 0.11 | 0.182 | 148 |
| Total meat production | **0.286** | **< 0.001** | 0.231 | 0.005 | 148 |
| Pork production | **0.484** | **< 0.001** | **0.303** | **< 0.001** | 148 |
| Beef and buffalo production | **0.164** | **0.046** | 0.086 | 0.297 | 148 |
| Sheep and goat production | -0.145 | 0.079 | -0.074 | 0.373 | 148 |
| Poultry production | **0.172** | **0.036** | 0.149 | 0.071 | 148 |
| Total meat consumption | **0.79** | **< 0.001** | **0.809** | **< 0.001** | 148 |
| Pork consumption | **0.714** | **< 0.001** | **0.564** | **< 0.001** | 148 |
| Beef and buffalo consumption | **0.551** | **< 0.001** | **0.583** | **< 0.001** | 148 |
| Sheep and goat consumption | 0.008 | 0.919 | 0.14 | 0.091 | 148 |
| Poultry consumption | **0.591** | **< 0.001** | **0.676** | **< 0.001** | 148 |

**Table S2.** Spearman correlation coefficients between ABU/kg meat and total ABU in livestock across six continents.

|  |  | Total ABU in livestock |  | N |
| --- | --- | --- | --- | --- |
| Items |  | r | p |  |
| ABU/kg meat in livestock | North America | **-0.516** | **0.012** | 23 |
|  | Oceania | -0.251 | 0.457 | 11 |
|  | Africa | 0.09 | 0.531 | 51 |
|  | South America | 0.483 | 0.112 | 12 |
|  | Europe | **0.469** | **0.002** | 40 |
|  | Asia | **0.551** | **< 0.001** | 44 |

**Table S3.** Spearman correlation coefficients between total meat production or consumption and total ABU or ABU/kg meat in livestock across six continents.

|  | Items | Total meat production |  | Total meat consumption |  | N |
| --- | --- | --- | --- | --- | --- | --- |
|  |  | r | p | r | p |  |
| Total ABU in livestock | North America | **0.984** | **< 0.001** | -0.332 | 0.122 | 23 |
|  | Oceania | **0.961** | **< 0.001** | -0.041 | 0.905 | 11 |
|  | Africa | **0.862** | **< 0.001** | -0.184 | 0.197 | 51 |
|  | South America | **0.958** | **< 0.001** | 0.503 | 0.095 | 12 |
|  | Europe | **0.872** | **< 0.001** | 0.198 | 0.221 | 40 |
|  | Asia | **0.876** | **< 0.001** | -0.227 | 0.138 | 44 |
| ABU/kg meat in livestock | North America | **-0.565** | **0.005** | 0.329 | 0.125 | 23 |
|  | Oceania | -0.318 | 0.34 | 0.373 | 0.259 | 11 |
|  | Africa | -0.093 | 0.515 | 0.142 | 0.319 | 51 |
|  | South America | 0.364 | 0.245 | 0.378 | 0.226 | 12 |
|  | Europe | 0.033 | 0.841 | -0.042 | 0.797 | 40 |
|  | Asia | **0.432** | **0.003** | -0.131 | 0.396 | 44 |

**Table S4.** Spearman correlation coefficients between GDP per capita or average LAYs and total ABU in livestock or in livestock, total meat production or consumption across five continents.

|  | Items | GDP per capita |  | Average LAYs |  | N |
| --- | --- | --- | --- | --- | --- | --- |
|  |  | r | p | r | p |  |
| Asia | Total ABU in livestock | -0.249 | 0.127 | 0.02 | 0.905 | 39 |
|  | ABU/kg meat in livestock | -0.294 | 0.069 | 0.083 | 0.617 | 39 |
|  | Total meat production | -0.152 | 0.357 | 0.102 | 0.536 | 39 |
|  | Total meat consumption | **0.814** | **< 0.001** | **0.772** | **< 0.001** | 39 |
| Europe | Total ABU in livestock | -0.069 | 0.673 | 0.087 | 0.592 | 40 |
|  | ABU/kg meat in livestock | **-0.654** | **< 0.001** | **-0.523** | **0.001** | 40 |
|  | Total meat production | 0.282 | 0.078 | **0.412** | **0.008** | 40 |
|  | Total meat consumption | 0.258 | 0.108 | 0.237 | 0.14 | 40 |
| South America | Total ABU in livestock | 0.5 | 0.207 | -0.238 | 0.57 | 8 |
|  | ABU/kg meat in livestock | **0.738** | **0.037** | 0.619 | 0.102 | 8 |
|  | Total meat production | 0.143 | 0.736 | -0.143 | 0.736 | 8 |
|  | Total meat consumption | **0.81** | **0.015** | -0.071 | 0.867 | 8 |
| Africa | Total ABU in livestock | -0.021 | 0.893 | -0.208 | 0.176 | 44 |
|  | ABU/kg meat in livestock | 0.195 | 0.205 | 0.077 | 0.621 | 44 |
|  | Total meat production | 0.11 | 0.478 | 0.007 | 0.964 | 44 |
|  | Total meat consumption | **0.469** | **0.001** | **0.354** | **0.018** | 44 |
| North America | Total ABU in livestock | 0.346 | 0.206 | 0.15 | 0.594 | 15 |
|  | ABU/kg meat in livestock | -0.193 | 0.491 | -0.039 | 0.889 | 15 |
|  | Total meat production | 0.414 | 0.125 | 0.2 | 0.475 | 15 |
|  | Total meat consumption | **0.746** | **0.001** | **0.682** | **0.005** | 15 |

**Table S5.** Spearman correlation coefficients between GDP per capita and Average LAYs.

|  |  | Average LAYs (years) |  | N |
| --- | --- | --- | --- | --- |
| Items |  | r | p |  |
| GDP per capita (US \$) | Globally | **0.854** | **p < 0.001** | 148 |
|  | Asia | **0.678** | **p < 0.001** | 39 |
|  | Europe | **0.651** | **p < 0.001** | 40 |
|  | South America | 0.262 | 0.531 | 8 |
|  | Africa | **0.545** | **p < 0.001** | 44 |
|  | North America | **0.732** | **0.002** | 15 |

**Table S6.**
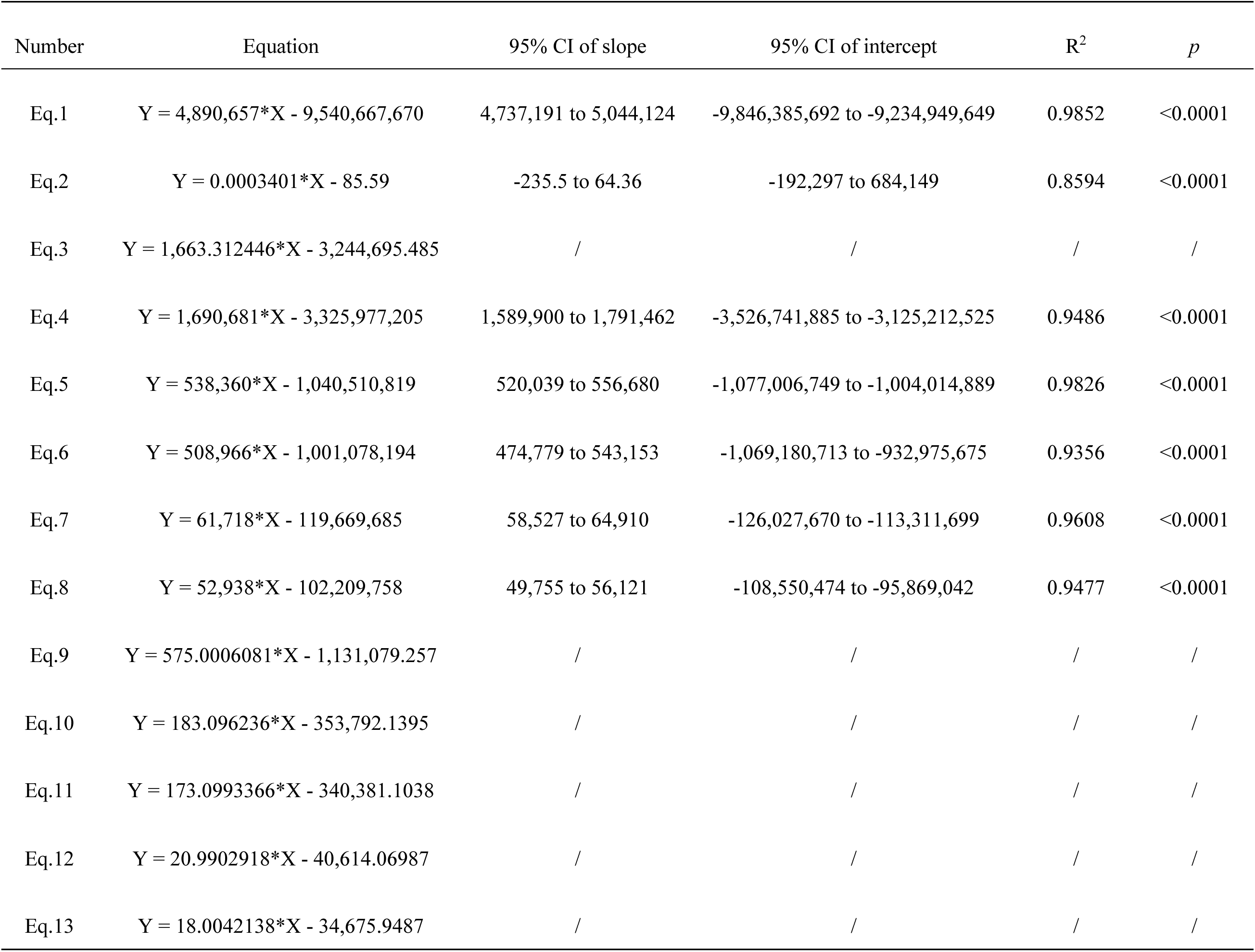
Table S6. Predictive equations and model parameters for total meat production, ABU/kg meat, and total ABU in livestock.

| Number | Equation | 95% CI of slope | 95% CI of intercept | R<sup>2</sup> | p |
| --- | --- | --- | --- | --- | --- |
| Eq.1 | Y = 4,890,657*X - 9,540,667,670 | 4,737,191 to 5,044,124 | -9,846,385,692 to -9,234,949,649 | 0.9852 | <0.0001 |
| Eq.2 | Y = 0.0003401*X - 85.59 | -235.5 to 64.36 | -192,297 to 684,149 | 0.8594 | <0.0001 |
| Eq.3 | Y = 1,663.312446*X - 3,244,695.485 | / | / | / | / |
| Eq.4 | Y = 1,690,681*X - 3,325,977,205 | 1,589,900 to 1,791,462 | -3,526,741,885 to -3,125,212,525 | 0.9486 | <0.0001 |
| Eq.5 | Y = 538,360*X - 1,040,510,819 | 520,039 to 556,680 | -1,077,006,749 to -1,004,014,889 | 0.9826 | <0.0001 |
| Eq.6 | Y = 508,966*X - 1,001,078,194 | 474,779 to 543,153 | -1,069,180,713 to -932,975,675 | 0.9356 | <0.0001 |
| Eq.7 | Y = 61,718*X - 119,669,685 | 58,527 to 64,910 | -126,027,670 to -113,311,699 | 0.9608 | <0.0001 |
| Eq.8 | Y = 52,938*X - 102,209,758 | 49,755 to 56,121 | -108,550,474 to -95,869,042 | 0.9477 | <0.0001 |
| Eq.9 | Y = 575.0006081*X - 1,131,079.257 | / | / | / | / |
| Eq.10 | Y = 183.096236*X - 353,792.1395 | / | / | / | / |
| Eq.11 | Y = 173.0993366*X - 340,381.1038 | / | / | / | / |
| Eq.12 | Y = 20.9902918*X - 40,614.06987 | / | / | / | / |
| Eq.13 | Y = 18.0042138*X - 34,675.9487 | / | / | / | / |

